# Subcellular assessment of algal carbon storage and chloroplasts across the microenvironmental landscape of a photosymbiotic coral

**DOI:** 10.1101/2025.08.19.671130

**Authors:** Andrea Catacora-Grundy, Netanel Kramer, Sofie Lindegaard Jakobsen, Michael Kühl, Johan Decelle, Daniel Wangpraseurt

## Abstract

Light availability plays a central role in shaping the photophysiology and energy metabolism of photosymbiotic organisms such as reef-building corals. Although light varies greatly within coral colonies, the effects of this spatial heterogeneity on the subcellular organization and energy storage of symbiotic algae (Symbiodiniaceae) remain poorly understood. Here, we combined microscale measurements of light and oxygen across both light-exposed upper regions and shaded basal regions of a *Favites abdita* colony with three-dimensional subcellular imaging using Focused Ion Beam Scanning Electron Microscopy (FIB-SEM). Our multi-scale approach revealed subcellular heterogeneity among symbiont populations, suggesting different cell cycle stages and physiological states across a spatial stratification in the coral. Subcellular morphometrics revealed that symbiont cells at the top of the colony were twice more voluminous than those at the shaded base with similar plastid volume occupancy. Compared to symbionts at the top of the colony, symbionts in the basal region accumulated nearly three times more starch relative to their cell volume. These findings show that light gradients within coral colonies shape symbiont morphology and energy storage patterns, with important implications for coral stress tolerance and resilience.

## Main body

Light is a pivotal parameter controlling photosynthesis and energy metabolism in multicellular organisms harboring symbiotic microalgae, such as reef-building corals [1]. These organisms host photosynthetic dinoflagellates within their tissues, forming a partnership where symbionts supply O_2_ and photosynthetically fixed carbon to the host, supporting growth and metabolism [2, 3], while the coral provides shelter, inorganic carbon and nutrients [1]. Despite extensive research on coral-symbiont biology, the influence of intra-colony light gradients on symbiont subcellular architecture and carbon dynamics remains poorly understood [4]. Symbiont populations are often treated as homogenous entities, despite dynamic light and chemical gradients within tissues that may structure the distribution and intracellular architecture of zooxanthellae *in hospite* [5, 6].

Light distribution in coral tissues varies across spatial scales, from colony-wide differences to microgradients within tissue [7–9]. Incident light can differ by up to 95% between exposed and shaded regions of branching corals, and only ∼10% of surface irradiance penetrates deeper tissue layers [7, 8]. Significant intra-colony variability in photosynthesis, carbon fixation, and photochemical efficiency has been documented [6, 10, 11], but subcellular responses of algal symbionts to light gradients, including changes in cell architecture, organelle size, and carbon storage, are largely unexplored [12]. To address this gap, we investigated the subcellular architecture of Symbiodiniaceae across light microhabitats within a massive *Favites abdita* colony at Heron Island, southern Great Barrier Reef. Fragments were collected from the directly light-exposed top and shaded base of the colony (Fig. 1a; Supplementary Information), corresponding to a ∼6-fold difference in scalar irradiance during midday sun exposure [7]. Using Focused-Ion Beam Scanning Electron Microscopy (FIB-SEM) and high-resolution microsensing, we analyzed changes in subcellular morphometrics, physiology, and the light microenvironment of Symbiodiniaceae. Microscale light and O_2_ concentration gradients were characterized with fiber-optic scalar irradiance and electrochemical O_2_ microsensors (∼50 µm tip diameter) [8]. Coral tissues were sampled *in situ*, fixed, and prepared for FIB-SEM imaging ([13]; Supplementary information). A 3D reconstruction pipeline quantified subcellular components, including cell volume, plastids, and starch granules [14]. These multiscale approaches enabled us to quantitatively link light microhabitats to the subcellular architecture and energy storage of the symbiotic dinoflagellates at different locations in the colony.

**Figure 1.**
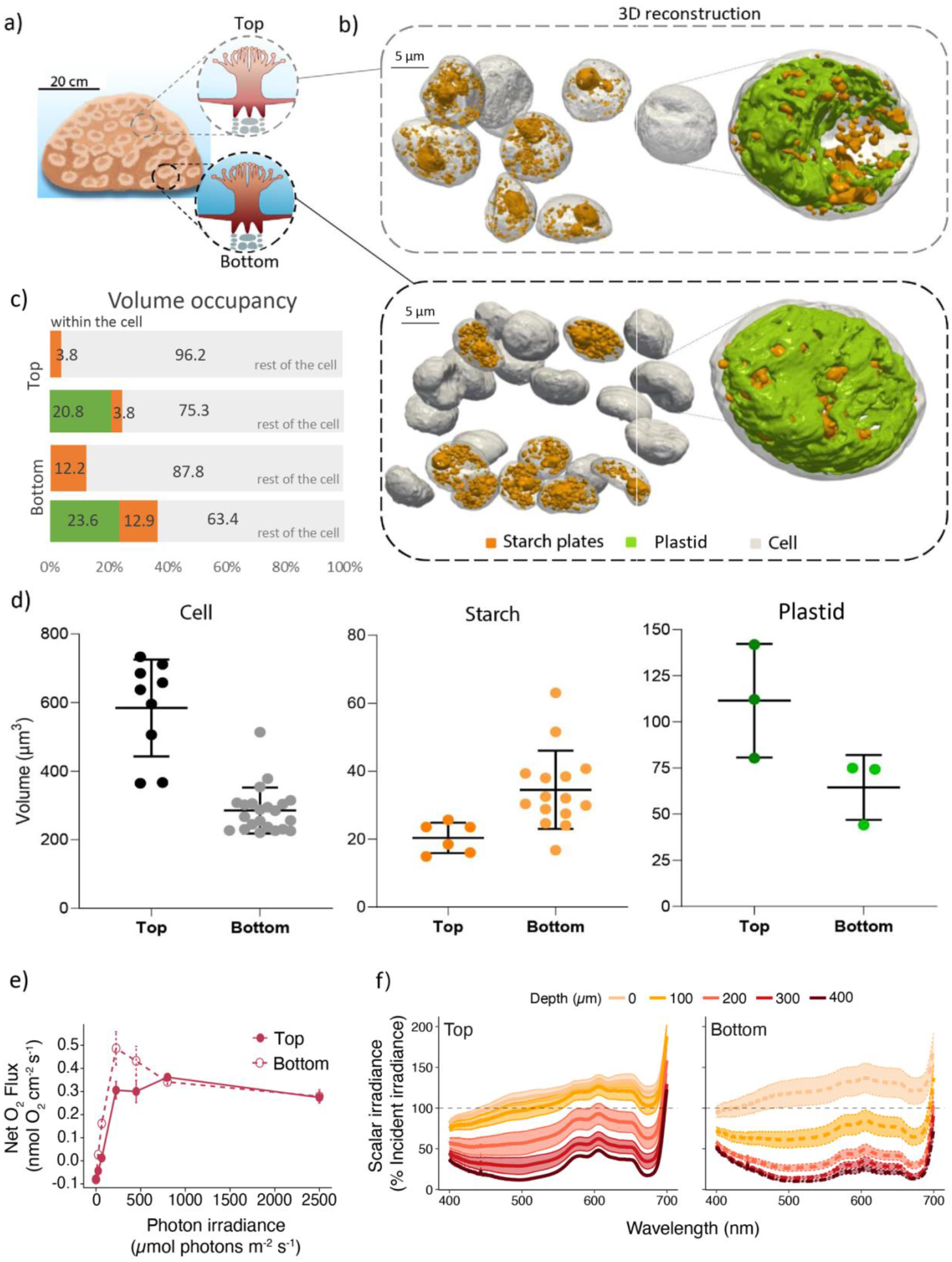
**(a)**Sketch of the massive coral *Favites abdita*, illustrating the top (light-exposed) and bottom (shaded) regions of the colony selected for sampling. **(b)** 3D FIB-SEM reconstruction showing the intracellular organization of symbiotic dinoflagellates within the endodermal tissue of *F. abdita*. **(c)** Starch granule volume occupancy (expressed as a percentage of total cell volume) in dinoflagellate symbionts from the top (light-exposed) and bottom (shaded) regions of the coral colony **(d)** Morphometric comparison of cell, starch granule, and plastids volumes (µm^3^) between the two regions, presented as mean ± SD. **(e)** Net O_2_ flux between coral coenosarc tissue and the overlaying water measured using O_2_ microsensors as a function of incident photon irradiance (data points represent means ± SD; *n* = 3-5 randomly chosen coenosarc tissue areas in the exposed and shaded parts of the coral colony, respectively). **(f)** Vertical microprofiles of spectral scalar irradiance in coenosarc tissue (solid lines represent mean spectra with SE is shown as colored areas; *n* = 5 random measurements in chosen tissue areas in the exposed and shaded parts of the coral colony, respectively). Scalar irradiances are normalized to incident downwelling spectral irradiance at the coral tissue surface.

Light microsensor measurements confirmed strong variation in internal light microclimates between colony regions (Fig. 1e-f). Light attenuation was strongest in shaded regions, consistent with their position, while high light-exposed top regions exhibited lower attenuation. These differences were reflected in net photosynthesis-irradiance curves, where bottom-region symbionts exhibited higher photosynthetic efficiency at low light, indicating low-light adaptation [15], while top-region symbionts exhibited greater maximum photosynthetic rates (Fig. 1e). Symbiont size and density also varied spatially in the host colony (Fig. 1a). While cells at the colony base were more densely packed (Fig. 1b), the cell volume of symbionts at the top were 2-fold higher (584.37 ± 140.80 µm^3^; n = 8) compared to cells at the base (285.04 ± 67.12 µm^3^; n = 22) (Fig. 1d). Although chloroplasts occupied a consistent relative volume (∼22%) (20.84 ± 2.6%, n=3 at the top and 23.63 ± 6.2%, n=3 in the bottom region – Suppl. Table 1) in microalga from both regions (Fig. 1c), their absolute volume was 1.7 times larger in symbionts from the upper colony’s region (111.52 ± 30.78 µm^3^; n = 3) than in the lower colony’s region (64.48 ± 17.61 µm^3^; n = 3) (Fig. 1d). This could suggest that symbionts modulate absolute plastid size in response to light availability without altering their overall cellular investment in the photosynthetic machinery.

Starch, the main form of carbon storage in Symbiodiniaceae, also varied spatially. Symbionts from the bottom region contained 1.7 times more starch granules (34.5 ± 11.5 µm^3^; n = 15) than those from the top (20.4 ± 4.5 µm^3^; n = 6). Relative to cell size, bottom-region symbionts had 3.2 times more starch (12.23 ± 4.35%; n=15; Fig. 1d) than top-region symbionts (3.80 ± 0.94%; n = 6; Fig. 1d). Thus, FIB-SEM quantification revealed distinct patterns of carbon storage between the light-exposed and shaded regions of the coral, while chloroplast proportions remained stable. Increased starch accumulation in shaded smaller symbionts suggests several non-exclusive scenarios: 1) less starch degradation, 2) higher carbon fixation and/or 3) reduced translocation of photosynthates to the host. Under low light, symbionts may prioritize starch accumulation as an energy reserve, whereas under high light, carbon is more rapidly translocated or used in other metabolic pathways. While low light typically induces larger chloroplasts to maximize light harvesting in microalgae like diatoms [14], the constant ∼22% chloroplast occupancy (∼22%) observed here suggests Symbiodiniaceae deploy a distinct photoacclimation strategy. The results align with studies showing light influences the balance of carbon storage and translocation in coral symbioses [16]: under nutrient-limited or low-light conditions, symbionts retain more fixed carbon as starch, likely as a survival strategy [17]. Our findings thus suggest that carbon allocation between symbiont and host varies over the coral colony in accordance with the local light conditions.

By revealing how intracellular symbionts adapt to light heterogeneity, our study highlights the need to view symbiont populations as physiologically heterogeneous, consistent with evidence of genetic and functional diversity within Symbiodiniaceae [18, 19]. Such heterogeneity may explain differential bleaching patterns within coral colonies, where shaded regions often retain symbionts longer during thermal stress [19]. The stratification of symbiont physiology documented here has important implications for coral stress responses. We speculate that the higher starch accumulation in shaded symbionts could reflect reduced carbon translocation to the host, but alternatively, larger energy reserves may render these symbionts more tolerant to bleaching. The latter interpretation aligns with field observations where shaded regions often retain symbionts longer during thermal stress, while exposed regions bleach first. However, such algal symbiont-centric view overlooks host metabolism and the heterogeneous distribution of holobiont-associated microbiomes across colonies [19, 20]. These intra-colony differences highlight the complex interplay between light environments, symbiont, microbiome and host physiology that ultimately determines coral health. Future research should integrate morphometric approaches with molecular profiling to map microbial diversity and gene expression patterns across light gradients, coupled with physiological measurements of carbon fixation and translocation rates, to elucidate the molecular mechanisms underlying the observed subcellular heterogeneities in zooxanthellae and their functional consequences for coral holobiont fitness.

## Acknowledgements

We thank the staff at Heron Island Research for excellent assistance during the field work. This research was conducted under a research permit from the Great Barrier Reef Marine Parks authority (G18-41571.1). We acknowledge funding by the Gordon and Betty Moore Foundation (GBMF9325 to D.W., GBMF9206 to M.K. & GBMF11532 to J.D.) and the US National Science Foundation (NSF-DBI # 2316391 and NSF-BSF # 2149925 to D.W.). A.C.G received funding from the LabEx GRAL (ANR-10-LABX-49-01), financed within the University Grenoble Alpes graduate school CBHEUR-GS (ANR-17-EURE-0003) Ecoles Universitaires de Recherche. J.D was supported by the ERC consolidator grant SymbiOCEAN (grant agreement no. 101088661).

