## Supplementary information for "Subcellular assessment of algal carbon storage and chloroplasts across the microenvironmental landscape of a photosymbiotic coral"

Measurements of *in situ* downwelling photon irradiance (400–700 nm) were conducted on a shallow reef flat adjacent to Heron Island Research Station (152°06′ E, 20°29′ S), Southern Great Barrier Reef, Australia. Measurements were performed between 9 AM and 6 PM (~sunset) on days characterized by clear skies and abundant sunlight, at water depths ranging from 0.3 to 2 meters (measured from the sediment benthos to the water surface) depending on tidal conditions. Low-tide measurements were conducted while snorkeling, and high-tide measurements were performed using SCUBA.

Data were collected using a hemispherical, underwater quantum irradiance sensor (LI-COR LI-192) connected to a LI-1400 data logger. The flat, light-diffusing disk of the sensor was positioned horizontally at the mean height of the coral colonies (~15 cm above the benthos) and approximately 2–3 meters from the colony of interest. In addition, ambient scalar irradiance of photosynthetically active radiation (PAR, 400–700 nm) was monitored using a miniature scalar irradiance sensor (3 mm diameter; Walz GmbH, Germany) connected to an underwater microsensor meter (UnderWater Meter system, Unisense A/S, Denmark). Net current flow velocity during selected study periods was estimated by tracking and timing the movement of small particles in the water column.

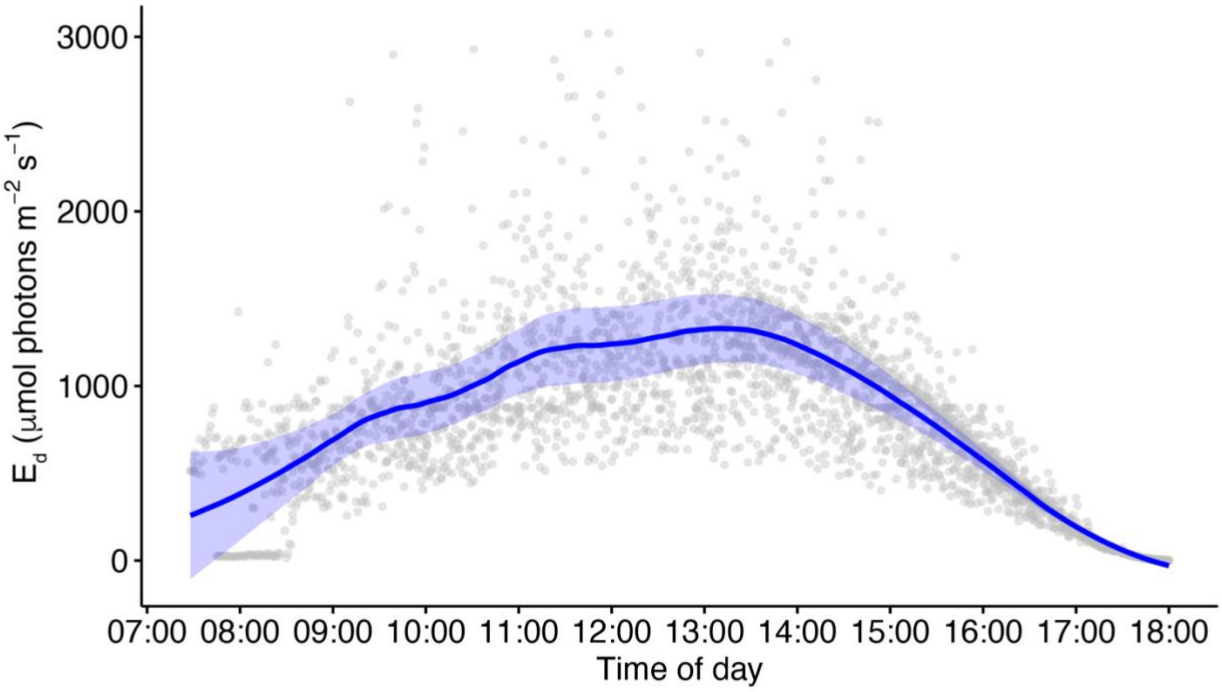


**Figure S1. Incident downwelling photon irradiance (400-700 nm) on a typical cloudless day on the shallow water reef flat near the Heron Island Research Station.** Trend line and shaded area are mean ± SE, respectively.

**Coral tissue sampling for FIB-SEM imaging and microsensing**

A single colony of the massive, thick-tissued coral *Favites abdita* was selected for analysis. This colony was located near the shoreline, growing on sediment with minimal algal cover at its base. Sampling was conducted around 4 PM using a hammer and chisel, collecting several small fragments (~1 cm thick, ~5 cm diameter) from both the top and bottom regions of the colony. Each fragment contained approximately 6–7 structurally intact polyps to assess potential differences in symbiont and tissue structure across the colony. Following collection, the samples were immediately transported in seawater to the outdoor aquaria at Heron Island Research Station (HIRS) and allowed to recover for 5 hours to minimize sampling stress. Samples designated for microsensor measurements were kept overnight in flow-through reef water before measurements began. For FIB-SEM preparation, tissue fixation was performed at approximately 10 using a fixative solution of 2.5% glutaraldehyde and 2.5% paraformaldehyde in 0.1 M phosphate buffer (pH 7.5), following Pernice et al. [1]. Fixation was carried out for 48 hours, after which the samples were transferred to PBS buffer for transport. Skeletal decalcification was then performed according to Pernice et al. [1] . Each fragment was cut into three smaller pieces using a bone cutter, stabilized in agarose within 50 mL containers, and immersed in ~20 mL of EDTA solution for decalcification, with the EDTA replaced every two days to ensure efficient decalcification.

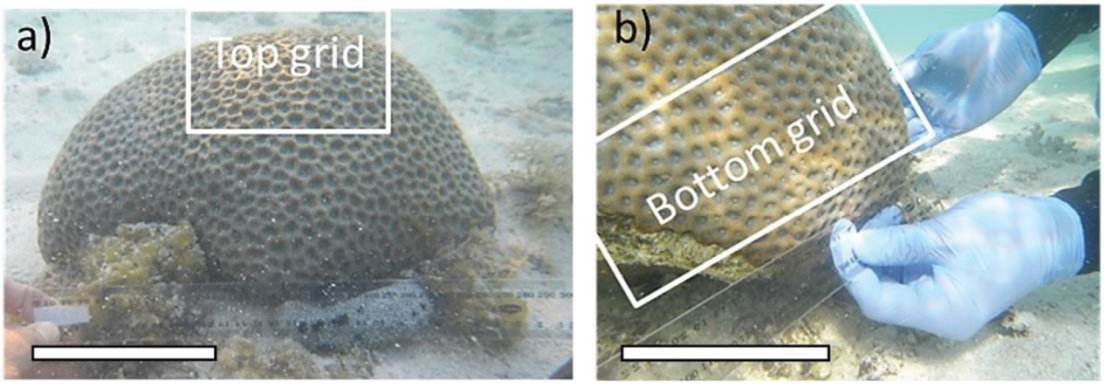


**Figure S2. Photographs of the *Favites abdita* coral investigated in this study show the top and bottom sampling areas. Scale bar = 10 cm.**

**FIB SEM imaging**

We applied FIB-SEM-based 3D reconstruction to understand the impact of light variation on the subcellular architecture and morphometrics of symbiotic dinoflagellates within coral host tissues. Our primary focus was on photosynthesis-related organelles (e.g., chloroplasts), cell volume, and carbon storage (e.g., starch). The following conditions were used for embedding the block for FIB-SEM acquisition: 2% $OsO_4$ buffer (1, 5 H), Wash MQ. (3 × 10–15 min), 1,2% uranyl acetate in water overnight, and washed with MQ. 3x 10-15 min. Dehydration:70% EtOH. 2x 15 min, 96% EtOH. (2x 15 min), 100% EtOH (3x 15 min), and 100% acetone (2 × 10–15 min). Epoxy: Epon/acetone 1:3 (45 min), 1:1 (45 min), 3:1 (45 min), Epon, 2-h or overnight and embedded at 60 °C for 24-h. Samples were trimmed on an ultramicrotome (Leica EM UC7), and trimmed sample blocks were mounted on ”stups” with PELCO® conductive silver paint and dried for minimum 3 hours. Samples were coated with Au (4 nm) using a sputter coater (Leica EM ACE200) and were imaged on a FIB-SEM system (FEI- Quanta 3D FEG) at the Core Facility for Integrated Microscopy, University of Copenhagen; https://cfim.ku.dk/).

Obtained image sequences were stacked, registered, and cropped using the Fiji software (https://imagej.net/Fiji) and its plugin Multistackreg. The transformation matrix was extracted from the plugin. The shearing caused by the 54° angle of the electron beam was corrected using a transformation matrix that was previously extracted using an in-house Python code (collab. P. Perrenot). Image processing was initiated with Fiji for cropping the selected cells and performing registration. Segmentation was performed using a semi-automatic method adopted from [2], based on pixel classification, using a 3D Slicer software (https://www.slicer.org/), and a supervised semi-automatic pixel classification mode was used to automatically segment 3 to 15 slices for each region of interest (ROI). Morphometric analyses were calculated using the Statistics Module in 3D Slicer. Three components of the symbiotic microalgae were segmented: the chloroplast, the starch granules in the cell, and the complete cell.

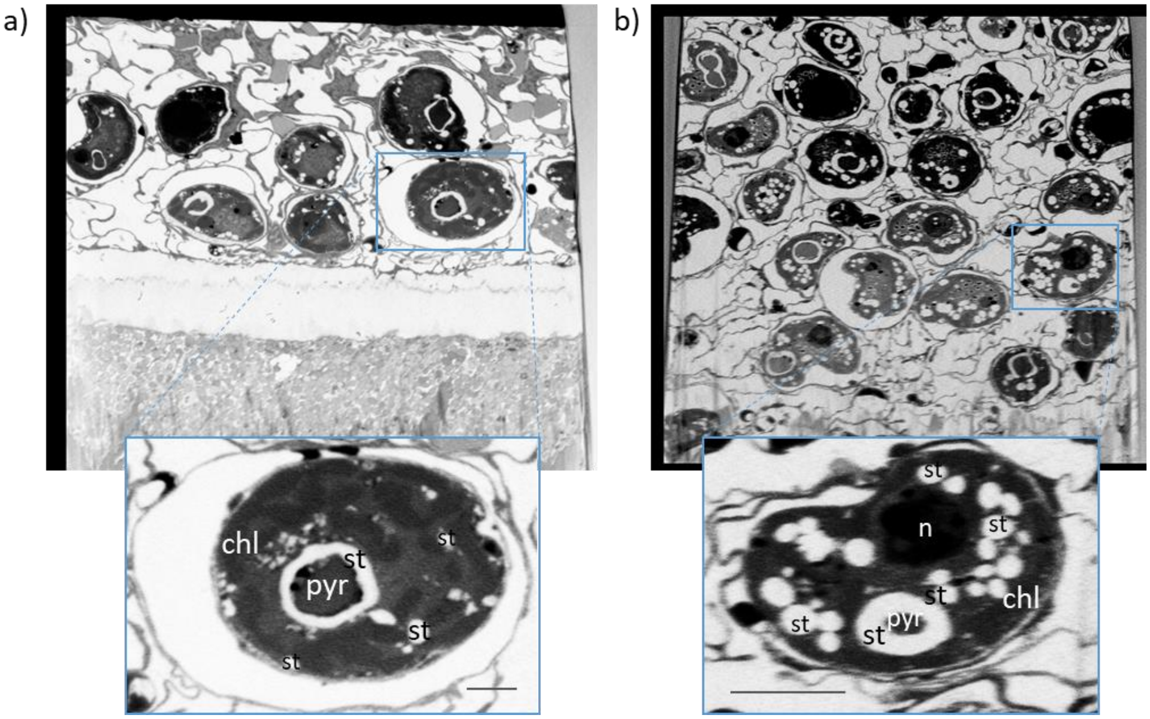


**Figure S3. Single FIB-SEM micrographs** of top **a)** and bottom. **b)** Sampling of symbiotic cells in the host *Favites abdita.* Zoom box in blue show some cellular structures of the cell: 'chl' = chloroplast, 'pyr' = pyrenoid, 'st' = starch, 'n' = nucleus. Scale bar = 2 µm.

### Coral tissue scalar irradiance and $O_2$ microsensing

Microscale scalar irradiance and $O_2$ microsensor measurements were performed according to established protocols [3, 4]. A fiber-optic scalar irradiance microsensor (tip diameter: 60 µm; https://www.zenzor.eu/)[2] connected to a spectrometer system (USB2000, Ocean Optics) was mounted on a motorized micromanipulator. The microsensor tip and the coral sample was observed under a dissection microscope to acquire precise vertical profiles within the coenosarc tissue of *Favites abdita*. Intratissue measurements were possible only in coenosarc tissue due to polyp retraction upon contact. For each fragment, three replicate vertical profiles were recorded to a depth of up to 400 µm, with additional profiles in thinner tissue regions to assess variability in attenuation coefficients. Downwelling photon irradiance (400–700 nm) was maintained at 300 µmol photons $m^{-2}$ $s^{-1}$ using a fiber-optic halogen light source (KL2500, Schott GmbH), as verified by measurements with a calibrated quantum irradiance sensor (LI-192 connected to a LI1400 meter; LI-COR).

Measurements of steady-state $O_2$ concentration profiles from the surrounding well-mixed seawater, across the diffusive boundary layer (DBL) and into the coral tissue were conducted under defined photon irradiance levels within ~5 minutes after stabilization under constant illumination using Clark-type $O_2$ microsensors (tip diameter: 25 µm, response time <0.5 s, stirring sensitivity ~1%; Unisense A/S, Denmark). Microsensors were calibrated in air-saturated and anoxic seawater, with $O_2$ concentrations calculated from percent air saturation using temperature- and salinity-corrected gas tables (Unisense, Denmark). The $O_2$ microsensors approached the coral surface at a ~10° angle relative to the vertically incident light. The net $O_2$ flux (J) between seawater and coral tissue across the DBL was calculated for each experimental light condition as [5]: $J=-D_0 \times dC/dz$, where $D_0$ is the molecular diffusion coefficient of $O_2$ in seawater at experimental salinity and temperature (Table available at Unisense.com, Denmark), and dC/dz is the linear $O_2$ concentration gradient measured in the DBL.
